# PTMlink: a database resource for co-variation of glycosylation and phosphorylation in human tumors

**DOI:** 10.64898/2026.11.23.768710

**Authors:** Effram Wei, Hongyi Liu, Yingwei Hu, Zhenyu Sun, Michael J. Betenbaugh, Hui Zhang

## Abstract

Protein glycosylation and phosphorylation can regulate each other, but the two protein modifications are usually studied separately, and their modification-specific co-variation across tumor types remains poorly characterized. PTMlink is a database of site-resolved co-variation between glycosylation and phosphorylation in the matched glycoproteomes and phosphoproteomes of 1,211 tumor samples from cohorts of the Clinical Proteomic Tumor Analysis Consortium, complemented by studies that perturbed glycosylation in human cells to examine its crosstalk with, and functional effects on, phosphorylation. Eligible glycosite–phosphosite pairs are scored within each tumor cohort by a covariate-adjusted partial rank correlation and combined across cohorts by meta-analysis, where specific glycoforms are scored in the same way and gene pairs are summarized. The strongest associations reproduce in held-out tumor types and show concordance in normal adjacent tissue, and the strongest pairs are enriched for known protein interactions and shared pathways; perturbation evidence recovers literature-documented relations that co-variation misses. Available through a website, an R package and a Model Context Protocol server, PTMlink is, to our knowledge, the first resource to provide a queryable score for site-specific relations between glycosylation and phosphorylation across tumor cohorts, with a framework that can extend to other protein modifications as matched measurements become available.

## 1 Introduction

Glycosylation and phosphorylation are among the most widespread protein modifications, and many cell-surface and secretory-pathway proteins, including receptor tyrosine kinases and integrins, carry both. A change in one can alter the other. Removing a single glycan from the epidermal growth factor receptor (EGFR) or from its relative ERBB3 (human epidermal growth factor receptor 3, HER3) lets the receptors oligomerize and become tyrosine-phosphorylated without ligand (1, 2). The glycosylation of adhesion and growth-factor receptors also shapes the kinase signaling that follows their engagement (3). Such relations have been established one protein at a time, largely in cultured cells. So far, the best-characterized crosstalk between glycosylation and phosphorylation involves intracellular *O*-GlcNAc (4).

Proteogenomic studies of human tumors make the coordination of the two modifications measurable, site by site, in clinical samples. The Clinical Proteomic Tumor Analysis Consortium (CPTAC) has profiled phosphopeptides in tumors of many cancer types (5–14). Intact glycopeptides were measured in the same samples, either by dedicated enrichment (13, 15, 16) or by re-analysis of the phosphoproteomic data (17, 18), and later cohorts extend the collection (19, 20). If specific glycosylation of one site and the phosphorylation of another are coupled, through a known or a yet unknown signaling pathway, their abundances can vary together across tumors. Co-variation that reproduces in independent tumor types and persists after accounting for shared protein abundance and tissue composition marks a candidate relation to guide experimental design and validation. Since co-variation has no direction, experiments that perturb glycosylation and measure phosphorylation in the same samples add a complementary causal view: how phosphorylation responds when glycosylation is altered (21–23).

Existing resources provide complementary views (Supplementary Table S1). Site databases such as Phospho-SitePlus, GlyGen, GlyConnect, iPTMnet and dbPTM catalog modification sites and their evidence (24–28). PTMcode infers associations between modifications from co-evolution, structural proximity and curation (29). These resources do not provide pooled glycosite-specific glycosylation and phosphosite co-variation estimates from matched tumor samples. Proteomic portals such as LinkedOmics and the Proteomic Data Commons serve per-cohort matrices and associations of one feature with all others, which LinkedOmics compares across cohorts (30, 31). A recent pan-cancer glycoproteomic resource correlates glycopeptides with phosphopeptides within a single cohort only (18). What remains unavailable is a pooled, site-pair query that reports the strength and consistency of an association across tumor types alongside evidence from glycosylation perturbations.

Here we describe PTMlink (https://ptmlink.biomarkercenter.org/), which meets this need for eligible glycosite-specific *N*-glycosylation–phosphosite pairs measured in the same human tumors. To our knowledge, it is the first resource to provide a queryable score for each such site-specific relation across cancer types, with each cohort’s estimate and, where it was measured, the normal adjacent tissue (NAT) estimate beside it. We show that its strongest pairs reproduce in held-out tumor cohorts and follow known protein relations, that it separates co-variation shared across tumor types from pairs that stand out in one, and that perturbation evidence extends it to documented relations that co-variation alone misses.

## 2 Materials and methods

### 2.1 Cohorts and data

PTMlink is built from tumor cohorts whose samples were profiled for both intact glycopeptides and phosphopeptides (Table 1): the ten cohorts of the CPTAC pan-cancer data freeze (Proteomic Data Commons release of 29 April 2021; the freeze cohorts), and new cohorts published since: gastric cancer (GC) and non-clear-cell renal cancers (Rare kidney cancers). The freeze cohorts use the harmonized phosphosite and protein abundances of the Baylor College of Medicine (32) and the intact-glycopeptide quantification of Johns Hopkins University; the later cohorts use the quantification tables of their source studies (Supplementary Methods S1).

**Table 1.**
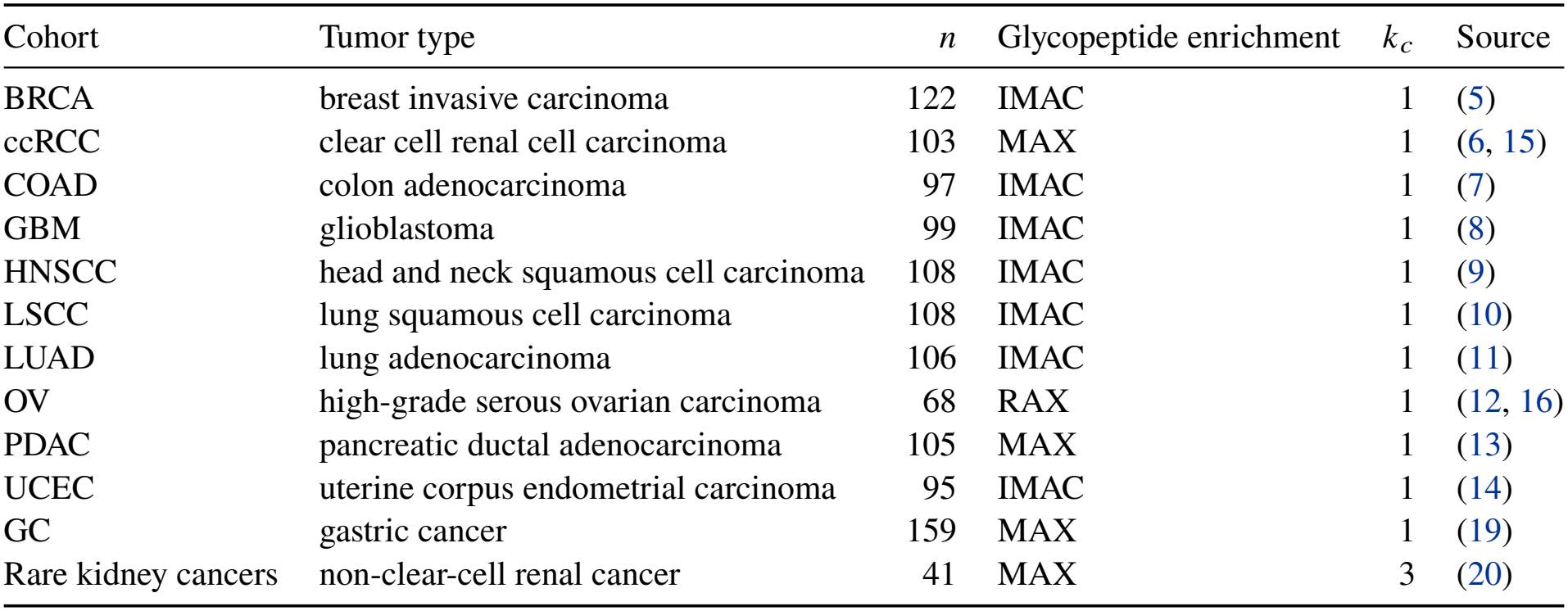
Tumor cohorts in PTMlink. *n*, number of tumors with both glycoproteome and phosphoproteome. Glycopeptide enrichment: MAX, mixed-mode anion exchange; RAX, Retain-AX anion exchange; IMAC, glycopeptides identified in the phosphopeptide-enriched data, as IMAC co-enriches sialylated glycopeptides. *k*_*c*_, covariates removed within the cohort.

In five cohorts, glycopeptides were enriched on a dedicated hydrophilic column. In the other seven they were identified in the phosphopeptide-enriched data, because immobilized metal affinity chromatography (IMAC) co-enriches sialylated glycopeptides (17, 18) from the same samples. Although these cohorts quantify fewer glycosites, their measured pairs contribute valid estimates of co-variation, and the held-out analysis evaluates each cohort separately.

Only tumors are scored, with one observation per patient (the median of repeated measurements). Missing values are not imputed, to avoid introducing artifacts into the data. NATs of six cohorts (ccRCC, COAD, HNSCC, LSCC, LUAD and PDAC) form separate cohorts that are displayed and compared but never pooled with tumors. All data are publicly released and de-identified, and no new human-subject data were generated.

### 2.2 Sites, glycoforms and identifiers

Every measurement is placed on a site key, so that cohorts processed with different software can be combined. A glycosite is a UniProt accession and the asparagine of an N-X-S/T sequon (X any amino acid except proline) in the canonical sequence (33). A phosphosite key retains the harmonized GENCODE v34 position of a phosphorylated serine, threonine or tyrosine; its mapped canonical UniProt position is reported separately and used in the stable identifier (Supplementary Methods S2). Gene symbols follow HGNC (34). A glycoform is a glycosite carrying one glycan composition. The 670 compositions are classed as paucimannose, oligomannose, glucosylated oligomannose, hybrid or complex, with fucosylation and sialylation recorded separately (Supplementary Methods S3).

### 2.3 Co-variation within cohorts

In cohort *c*, a glycoform–phosphosite pair is scored over the *N*_*c*_ tumors, at least 20, in which the glycoform, the phosphosite and every covariate are quantified. The partial Spearman correlation *ρ*_*c*_ is the correlation of the two ranked abundances after least-squares removal of the ranked covariates. The covariate is tumor purity (the ESTIMATE score (35)). The rare kidney cohort mixes histological types of non-clear-cell renal cancer; it uses the purity estimate published with the study and two indicators of histological group (papillary and other types, against oncocytic tumors). The number of covariates, *k*_*c*_ (Table 1), reduces the cohort’s degrees of freedom and its weight. Every correlation is computed within one cohort, so differences between cohorts in enrichment, quantification or measurement depth do not enter any correlation. Benjamini–Hochberg (BH) correction (36) over a cohort’s tests gives the cohort’s own *q* value.

A glycosite’s statistic in a cohort is that of its representative glycoform, the one quantified with the phosphosite in the most tumors. The representative glycoform may differ between cohorts, and it summarizes the best-covered glycoform rather than total site occupancy. Glycoform-level estimates are pooled only across cohorts that measure the same glycoform.

### 2.4 Pooling across cohorts

Cohorts, each processed separately, are combined by a fixed-effect meta-analysis on the Fisher scale (37):

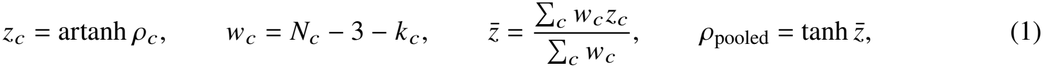

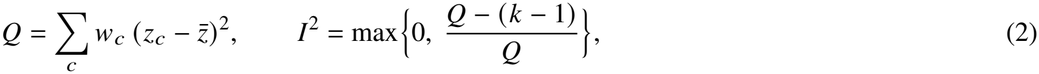

where *k* is the number of cohorts measuring the pair. Significance comes from a weighted Stouffer combination of the cohorts’ Student tests. Evidence, the ranking score, is log_10_ of the two-sided *P* value of this pooled test (Supplementary Methods S10). The null hypothesis is that the pair co-varies in none of its cohorts, so heterogeneity is reported beside each pair rather than absorbed into the test: *Q, I*^2^ (undefined at *k* = 1) and every cohort’s estimate. BH correction over every scored site pair gives its *q* value, and pairs at *q* < 0.05 are called significant (an evidence of 2.35 or more in this release); glycoform pairs form a separate family. A gene pair is represented by its strongest site pair. It is ranked by the percentile of that evidence among gene pairs with a similar number of site pairs, because the strongest of many site pairs is large by chance alone. In simulations and in permutations of measured pairs without co-variation, the pooled test rejected a single pair at close to its nominal rate at every number of cohorts examined (Supplementary Figure S1 and Supplementary Methods S10).

### 2.5 Perturbation evidence

Three studies perturbed *N*-glycosylation and measured phosphorylation in the same human cells: A498 renal carcinoma cells treated with eight doses of tunicamycin, an *N*-glycosylation inhibitor, and a vehicle control, with and without epidermal growth factor (EGF) (21); HEK293 cells carrying knock-ins and knock-outs of *N*-glycan biosynthesis genes (22); and A498 cells treated with sialidase, which removes sialic acids, with and without EGF (23). A site pair carries a study’s evidence when a glycoform of its glycosite and its phosphosite both responded to the perturbation, or when the dose-response study observed the pair itself as a correlation across doses (Supplementary Methods S4). A site pair is scored when it is measured in at least two tumor cohorts or carries perturbation evidence; perturbation evidence admits pairs but never enters the score. The database reports the significant site pairs and those with perturbation evidence.

### 2.6 Annotations

Two influences are annotated rather than removed: abundance shared by the two parent proteins, and tissue composition beyond purity. Removing them would also remove signal, because *N*-glycosylation governs the folding, trafficking and turnover of its carrier protein (38) and the composition scores are built from protein abundances of the same samples. Each pair carries the annotations of Table 2.

**Table 2.** Annotations attached to each reported pair. None changes the ranking.

| Annotation | Definition |
|---|---|
| Parent-protein correlation | the pooled correlation of the gene-level abundances of the two parent proteins |
| Site–protein coupling | the larger absolute median correlation of either site with the abundance of its own protein |
| Tissue-program labels | whether either site tracks tumor purity or a tissue-composition or secretory-pathway score |
| Program-adjusted correlation | *ρ* after removing purity, the immune–stromal contrast and the plasma-protein, proliferation and myeloid scores within each cohort, with its 95 % interval |
| Leave-one-cohort-out | the smallest evidence and the range of *ρ* over single-cohort removals, and the cohort whose removal moves *ρ* most |
| Tumor specificity | whether a cohort's *ρ* differs from that of the other cohorts pooled (*z*<sub>Δ</sub>, *q*<sub>Δ</sub>; see Tumor specificity in Materials and methods) |
| Perturbation evidence | the study that supports the pair and whether it observed the exact pair |

The parent-protein correlation and the program-adjusted correlation are pooled as the site pairs are. A tissue-program label marks a phosphosite, or a glycoform of a glycosite, whose median correlation across cohorts with purity or with one of eleven tissue-composition and secretory-pathway scores (Supplementary Methods S5) reaches |*ρ*| ≥ 0.6. A label shows that a site follows a score; the program-adjusted correlation shows whether the pair’s co-variation remains once the main scores are removed. Its interval is an annotation and carries no multiplicity correction.

A candidate screen combines the annotations. It keeps significant cross-protein pairs measured in at least six cohorts that meet four further conditions. First, the parent proteins barely co-vary (|*ρ*| < 0.15). Second, neither site follows its own protein (site–protein coupling below 0.6) or carries a tissue-program label. Third, the program-adjusted interval excludes zero. Fourth, the evidence is robust to the removal of any single cohort: the smallest share of evidence retained has a percentile above 0.5 among reported pairs with the same number of cohorts. A pair with a missing value for any numerical filter is excluded.

### 2.7 Reproducibility and known relations

#### Held-out cohorts

Each cohort in turn was held out and the others pooled. Among the cross-protein site pairs measured in at least two of the pooled cohorts, the 1,000 with the strongest pooled evidence were selected. Their correlations in the held-out cohort were regressed on their pooled correlations, and the share of pairs with the pooled sign was recorded. A slope of one means that the held-out cohort reproduces the pooled correlations at full strength, and a slope of zero that the two are unrelated (Supplementary Methods S6). The selected pairs were counted as reproduced in a cohort when at least 100 of them were measured in it and the lower 95 % bootstrap bounds exceeded zero for the slope and one half for the share with the pooled sign.

#### Normal adjacent tissue

The six NAT cohorts with at least 40 samples were scored as the tumors were, without a purity covariate, and pooled. Concordance was measured in two ways for cross-protein pairs significant in tumors and measured in at least two NAT cohorts: the share of pairs whose NAT correlation has the tumor sign, and the Spearman correlation between the tumor and NAT statistics. Each was compared with non-significant pairs with the same distribution of cohort counts (Supplementary Methods S6).

#### Known interactions and pathways

Cross-protein gene pairs among the reported pairs were tested, by evidence bin, for a STRING v12.0 experimental or curated-database edge (39) and for a shared Reactome pathway of at most 200 genes (40), against a random sample of all scored pairs. Since co-abundant and well-studied proteins have more known relations, the comparison was stratified by the parent proteins’ correlation and by each gene’s degree in the interaction and text-mining channels of STRING and combined as a Mantel–Haenszel risk ratio (41, 42) (Supplementary Methods S7).

### 2.8 Literature relations

Literature relations are published experiments in which altering the *N*-glycosylation of a named glycoprotein changed a phosphorylation on a named protein. Each cross-protein relation was compared with up to ten matched gene pairs. A matched pair shares the relation’s phosphoprotein, so that a frequently studied readout is its own control, and its glycoprotein resembles the relation’s in perturbation coverage, literature attention and measurement depth (Supplementary Methods S8). A relation has perturbation-only support when a site pair of its two proteins carries perturbation evidence and none is significant in the tumors. The statistic is the number of relations with this support divided by the number expected from their matched pairs (one at chance), with intervals from resampling glycoproteins. The list of relations was fixed before the twelve-cohort analysis.

### 2.9 Tumor specificity

Each cohort’s correlation for a pair was compared with the pooled correlation of the other cohorts that measure it by a *Z* statistic, *Z*_Δ_, whose variance combines both (Supplementary Methods S9); BH correction within the cohort gives *q*_Δ_. A pair stands out in a cohort when it is significant in the cohort’s own test, *q*_Δ_ < 0.05 and its correlation there is stronger than in the other cohorts. It is cohort-specific when it stands out in one or two cohorts, and it belongs to the shared background when it is measured in at least three cohorts, significant when pooled and stands out in none. The same section reports two checks of the statistic: between random halves of each cohort, and between two phosphoproteomic pipelines applied to the same tumors. Since *z*_Δ_ grows with the cohort’s weight, cohorts were also compared at equal size: each cohort’s correlations were recomputed on 30 randomly drawn tumors per pair, ten times.

## 3. Results

### 3.1 A site-resolved map in which evidence accumulates across tumor types

PTMlink scores 74,510,821 glycosite–phosphosite pairs that link 1,099 glycoproteins to 8,447 phosphoproteins, together with 412,549,021 glycoform–phosphosite pairs and 6,759,705 gene pairs (Figure 1A). Of the site pairs, 6,643,986 (8.92 %) are significant and 499,533 carry perturbation evidence; together they form 7,094,597 reported pairs on 2,521 glycosites and 54,523 phosphosites of 9,020 genes. Coverage differs between cohorts, from 84.4 % of reported pairs measured in GC to 10.0 % in OV (Figure 1B). The share of pairs that reach significance rises with the number of cohorts that measure them, from 5.2 % of pairs measured in two cohorts to 23.9 % of those measured in all twelve (Figure 1C): evidence accumulates as more cohorts measure a pair.

**Figure 1.**
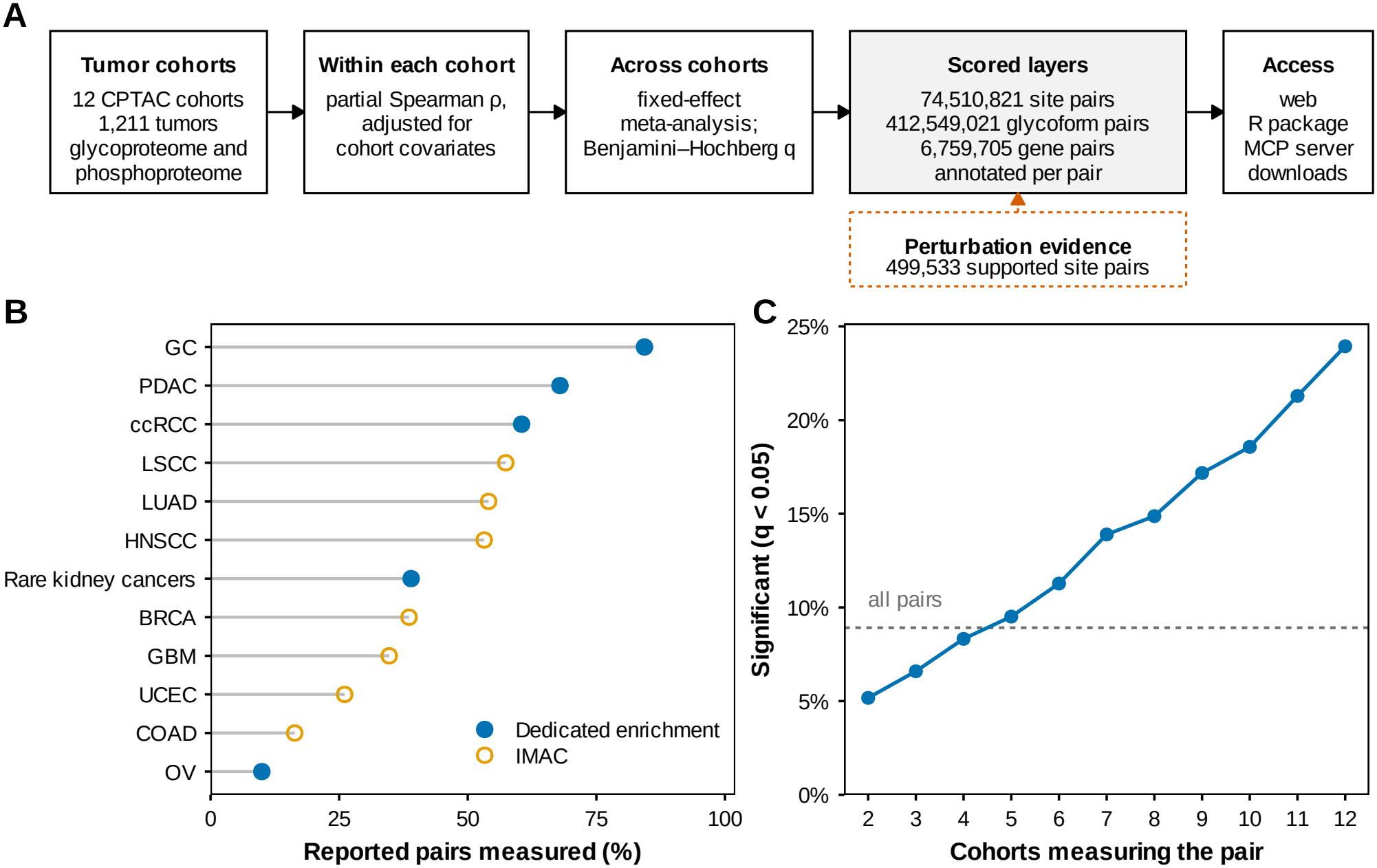
The PTMlink resource. (**A**) Workflow: glycoform–phosphosite pairs are scored within each tumor cohort, pooled across cohorts and summarized to gene pairs; perturbation evidence annotates pairs without entering the score. (**B**) Share of reported pairs measured in each cohort. Filled, dedicated glycopeptide enrichment; open, IMAC. (**C**) Share of site pairs significant at *q* < 0.05 by the number of cohorts measuring the pair; dashed line, all scored site pairs.

### 3.2 Co-variation reproduces across tumor types

Each cohort was held out in turn, and the 1,000 strongest pairs were selected from the other cohorts. In twelve of twelve held-out cohorts, the selected pairs reproduced, with either type of glycopeptide enrichment. The slope of held-out on pooled correlations was 0.35 to 1.20, and between 78.9 % and 98.7 % of the pairs had the pooled sign (Figure 2A).

**Figure 2.**
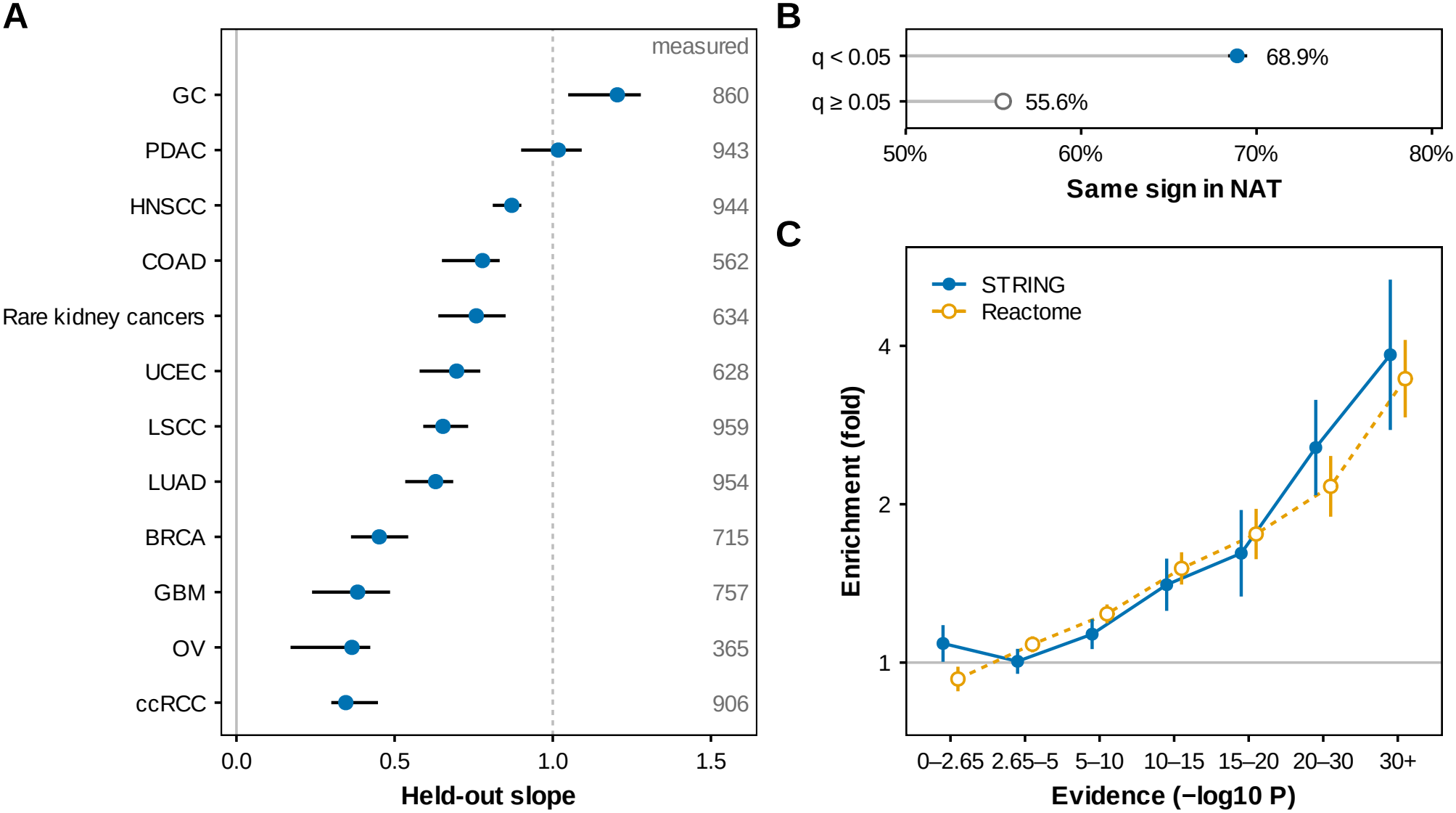
Co-variation reproduces across tumor types and follows known protein relations. (**A**) Slope of held-out on pooled correlations for the 1,000 strongest pairs selected without each held-out cohort; bars, 95 % bootstrap intervals (hidden where narrower than the point); numbers, selected pairs measured in the held-out cohort; line at one, full-strength reproduction. (**B**) Share of pairs whose pooled correlation in normal adjacent tissue (NAT) has the tumor sign, for pairs significant in tumors (filled) and non-significant pairs (open); bars, 95 % bootstrap intervals. (**C**) Enrichment for STRING experimental or database edges (filled, solid line) and Reactome co-membership (open, dashed line) by bin of evidence, relative to a random sample of scored gene pairs; bars, 95 % intervals.

Tumor co-variation was also concordant in NAT (Figure 2B). Of 4,510,182 pairs significant in tumors and measured in at least two of the six NAT cohorts, 68.9 % (95 % CI 68.4 %–69.5 %) had a pooled NAT correlation of the tumor sign, against 55.6 % of non-significant pairs. The tumor and NAT statistics were correlated (Spearman 0.445, 95 % CI 0.432–0.459, against 0.167). The concordance is partial, as expected for a tissue of different composition. Tumors and NAT share patients and isotope-labeling sets, so the significant pairs are compared with non-significant pairs measured in the same samples.

### 3.3 Co-variation follows known protein relations

The strongest pairs were enriched for known interactions and shared pathways (Figure 2C). Gene pairs with evidence above 30 were 3.8-fold (95 % CI 2.8–5.3) enriched for a STRING experimental or database edge and 3.5-fold (2.9–4.1) for Reactome co-membership. Both folds are estimated within strata of the parent proteins’ correlation and of each gene’s interaction and text-mining degree, so the enrichment persists after stratification by shared protein abundance and research attention (Supplementary Methods S7; by decile of parent-protein co-variation, Supplementary Figure S2B).

Supplementary Table S2 places six examples involving published protein relations beside six untested candidates. Two of the published relations illustrate how PTMlink presents a pair. Integrin *β*4 (ITGB4) N327 co-varies with AKT1 S124 over four cohorts (*ρ* = 0.38, evidence 8.87), consistent with reports that removing the *N*-glycans of integrin *β*4 reduces Akt activation (43, 44). The activating sites of AKT1, T308 and S473, are not among the reported sites, and the role of S124 in this relation remains to be determined. EGFR N603 co-varies with ALG3 S13 in the same direction in all five cohorts that measure it (*ρ* = 0.22, evidence 5.18), although the parent proteins barely co-vary, and the pair carries perturbation evidence. AKT phosphorylates ALG3 at Ser11 and Ser13, and this phosphorylation is required for the *N*-glycosylation of EGFR and other cell-surface receptors (45); here phosphorylation acts on glycosylation. Additional examples involve integrin α*V*–AKT, integrin *β*1–paxillin, PD-L1–STAT1 and CD44–ERK, for which glycosylation perturbations alter downstream phosphorylation (46–49). The table illustrates these protein relations at measured sites; the published experiments do not establish regulation between each listed residue pair.

### 3.4 Pairs that stand out in individual tumor types are separated from the shared background

The number of pairs that stand out in a cohort depends on how many tumors measure each pair, so cohorts were compared with every pair tested on 30 randomly drawn tumors (Materials and methods). Cohorts still differed (Figure 3A): the share of tested pairs that stood out was 0.71 % in GC, 31 times that of the median cohort (0.023 %). This met the criterion set before the test. PDAC had the second-largest share (0.51 %).

**Figure 3.**
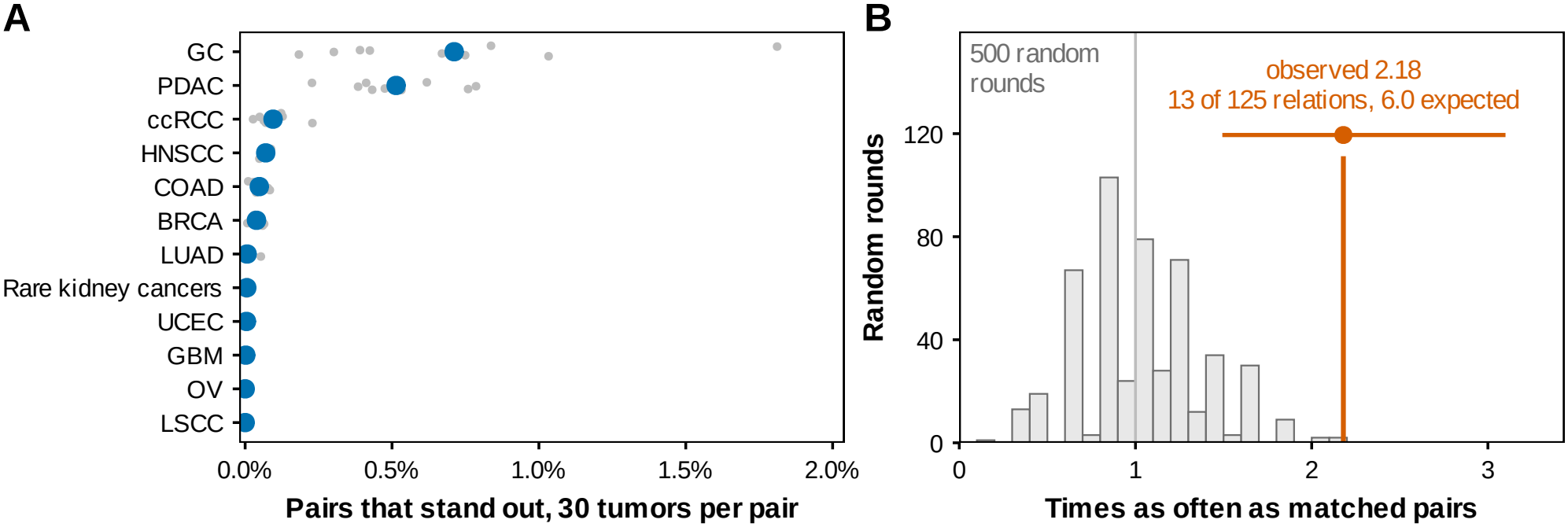
Beyond the shared background. (**A**) Share of the pairs tested in each cohort that stand out from the other cohorts pooled, with every pair tested on 30 randomly drawn tumors; small points, ten random draws; large points, their mean. (**B**) Literature relations with perturbation support and no significant tumor co-variation, as a ratio to the number expected from matched gene pairs. Histogram, 500 rounds in which every relation was replaced by one of its matched pairs; gray line, a ratio of one; vermillion line and bar, the observed ratio and its 95 % bootstrap interval.

### 3.5 Perturbation evidence reaches documented relations that co-variation misses

Of the literature relations, 125 cross-protein relations on 43 glycoproteins could be matched in PTMlink. Relations with perturbation support but without significant tumor co-variation occurred 2.18 times as often as among their matched pairs (95 % CI 1.49–3.10): 13 relations against 6.0 expected (Supplementary Table S3). In 500 rounds in which every relation was replaced by one of its own matched pairs, the ratio never reached this value (Figure 3B). None of the 13 relations was established with tunicamycin-class inhibitors, the drug of the dose-response study, so the excess does not arise because the literature and the perturbation evidence used the same drug.

### 3.6 The annotations narrow co-variation to testable hypotheses

The annotations can be combined into a screen for candidates (Materials and methods). The screen retains 256,097 pairs, 2,654 of them with perturbation evidence (Supplementary Table S14; Supplementary Table S2 lists six of them). One of these, gp130 (IL6ST) N131 × TMX1 S270, shows how an untested candidate is read. The two sites co-vary in the same direction in all seven cohorts that measure them (469 tumors; *ρ* = 0.23, evidence 5.90, *I*^2^ = 0.15), and the dose-response study observed the pair itself. The parent proteins do not co-vary, and the co-variation persists after adjustment for tissue programs. In NAT, by contrast, the pair’s correlation is near zero or negative in every cohort that measures it; whether the relation is confined to tumors remains to be tested. N131 is glycosylated in the human gp130 extracellular domain (50), and blocking *N*-glycosylation inhibits the activation of JAK and STAT3 by an IL-6-family cytokine in rat cardiomyocytes (51). TMX1 is a transmembrane oxidoreductase of the endoplasmic reticulum that acts preferentially on cysteine-containing membrane proteins (52), a class that includes gp130; S270 lies in its cytosolic tail (further background in Supplementary Table S2). The pair suggests a testable link between the glycosylation of a cytokine receptor and the phosphorylation of an oxidoreductase that acts on membrane proteins of its class.

### 3.7 Access

The website (https://ptmlink.biomarkercenter.org/) is a static application that requires no database server. It accepts a gene symbol, alias, UniProt accession or site. A gene page ranks the partner sites by evidence, with filters that state how many pairs they hide. A pair page shows every annotation, each cohort’s estimate beside the other cohorts pooled and NAT, and every glycoform of the glycosite. Network, tumor-specificity, list and status views complete the website, and every table exported from it carries a provenance header.

The `ptmlink` R package and a Model Context Protocol (MCP) server for AI agents read the reported site-pair files. Both apply the default view of the website: the significant cross-protein pairs measured in at least two cohorts, with the number of pairs each filter hides. Both provide the rules for reading a pair and identify every pair by a stable identifier built from UniProt accessions and residue positions, such as PTMG:P40189:N131|PTMP:Q9H3N1:S270; NAT and cohort-against-background comparisons are website features. Software versions and the version record of release v1.0.0 are in Supplementary Methods S11.

## 4 Discussion

PTMlink makes co-variation between *N*-glycosylation and phosphorylation measurable site by site in human tissue. The strongest pairs reproduce in held-out tumor types, show concordance in NAT and are enriched for known interactions and shared pathways. These results indicate a component of co-variation that is shared across the measured tissues and follows functional relations between proteins.

Tumor co-variation and experimental perturbation measure different concepts: which modified-peptide abundances move together at steady state across many individuals, and how phosphorylation changes when glycosylation is deliberately altered in one cell system. Literature relations carry perturbation support without tumor co-variation more often than matched gene pairs, which suggests that some glycosylation-dependent signaling occurs under conditions that natural variation between tumors does not sample. Conversely, co-variation is symmetric, so it can capture regulation that runs from phosphorylation to glycosylation, such as the AKT-dependent phosphorylation of ALG3 that EGFR glycosylation requires (45). Showing both kinds of evidence side by side lets a user distinguish a pair that co-varies in patient tumors from one that responds only when glycosylation is altered.

Pairs that stand out in a single tumor type are candidates for tissue-specific regulation. They concentrate in gastric and pancreatic cancer even when every cohort tests each pair on the same number of tumors, so cohort size alone does not explain the concentration. Both tumor types carry abundant, variable stroma, so a cohort-specific pair can reflect tissue composition as well as tumor biology. PTMlink therefore shows each cohort’s estimate beside the other cohorts pooled.

PTMlink complements other available post-translational modification (PTM) databases (Supplementary Table S1): it adds co-variation measured in human tissue, per-cohort effects, glycoform resolution and explicit annotation of the confounders that a correlation cannot exclude. The candidate table deposited with this article lists the pairs that pass the complete screen.

Three properties of the data limit what a pair can show. First, a correlation is observational and has no direction, and perturbation evidence shows only that a pair’s sites respond when glycosylation is altered; each pair is a hypothesis for a targeted experiment. Second, shared protein abundance and tissue composition can move two sites together without any regulation between them; the annotations expose these explanations, and the candidate screen removes every labeled pair. Third, coverage differs several-fold between cohorts and favors abundant secreted and extracellular-matrix glycoproteins, so the absence of a pair does not show the absence of a relation. Because each correlation is computed within one cohort, these differences affect which pairs are measured, not the correlations themselves.

Each archived release carries stable pair identifiers, an immutable build record and file checksums, so later releases can add tumor cohorts, perturbation studies and a unified re-analysis of the raw CPTAC data without breaking citations. The same framework extends to other PTMs as matched site-level data become available.

## Acknowledgements

We thank the CPTAC investigators, study participants and the authors of the source studies for generating and sharing the data used in PTMlink. We acknowledge the NCI Data Jamboree for providing technical support and data resource expertise.

AI tools, including Claude (Anthropic), assisted literature curation and text mining for literature relations, figure formatting, and language editing. The authors reviewed, verified and revised all AI-assisted content and take responsibility for the final work.

## Author contributions

Effram Wei: Conceptualization, Methodology, Software, Formal analysis, Data curation, Validation, Visualization, Writing—original draft, Writing—review & editing. Hongyi Liu: Conceptualization, Investigation, Data curation. Yingwei Hu: Investigation, Data curation, Software (website development and deployment). Zhenyu Sun: Investigation, Data curation. Michael J. Betenbaugh: Supervision, Writing—review & editing. Hui Zhang: Conceptualization, Supervision, Project administration, Funding acquisition, Resources, Writing—review & editing.

## Supplementary Information

Supplementary Methods S1–S11, Supplementary Tables S1–S15 and Supplementary Figures S1–S2 accompany this article. Supplementary Tables S4–S15 are provided as one Excel file.

## Conflict of interest

The authors declare no conflict of interest.

## Funding

This work was supported by the National Cancer Institute of the National Institutes of Health through the Clinical Proteomic Tumor Analysis Consortium (CPTAC, U24CA271079 to H.Z.), the Early Detection Research Network (EDRN, U2CCA271895 to H.Z.), the Pancreatic Cancer Detection Consortium (PCDC, U01CA274514 to H.Z.), and by the Johns Hopkins Brady Urological Institute Kidney Cancer Program Pilot Research Grant (to H.L.). This manuscript is the result of funding in whole or in part by the National Institutes of Health (NIH). It is subject to the NIH Public Access Policy. Through acceptance of this federal funding, NIH has been given a right to make this manuscript publicly available in PubMed Central upon the Official Date of Publication, as defined by NIH.

## Data availability

The PTMlink website (https://ptmlink.biomarkercenter.org/) is publicly available and serves release v1.0.0: each reported pair with its per-cohort estimates, annotations and glycoforms, the tumor-specificity and NAT comparisons, and an export of each result table with a provenance header. The ptmlink R package and the MCP server read the same reported site-pair files and are openly available at https://github.com/Effram360/PTMlink; the release is archived on Zenodo with a DOI on acceptance. Code is licensed under MIT and PTMlink data under CC BY 4.0, subject to the source-data terms. The complete candidate screen and the script that reproduces it are deposited on Zenodo, with access for the reviewers until publication; the full data release follows publication of the article. The tumor matrices are publicly released CPTAC data from the Proteomic Data Commons (31) and the quantification tables of the cohorts published later (Table 1). The perturbation evidence was built from the processed tables of the three cited studies (21–23). Those studies deposited their mass-spectrometry data in PRIDE: PXD071440 (dose response), PXD071742 (gene editing) and PXD078365 (sialidase, released with its study). Reference resources, namely STRING v12.0 (39), Reactome (40), UniProt (33), HGNC (34), iPTMnet (27), PTMcode v2 (29) and NCBI gene2pubmed, are pinned by version, retrieval date or checksum in the release’s version record.

## Table of Contents (TOC) figure

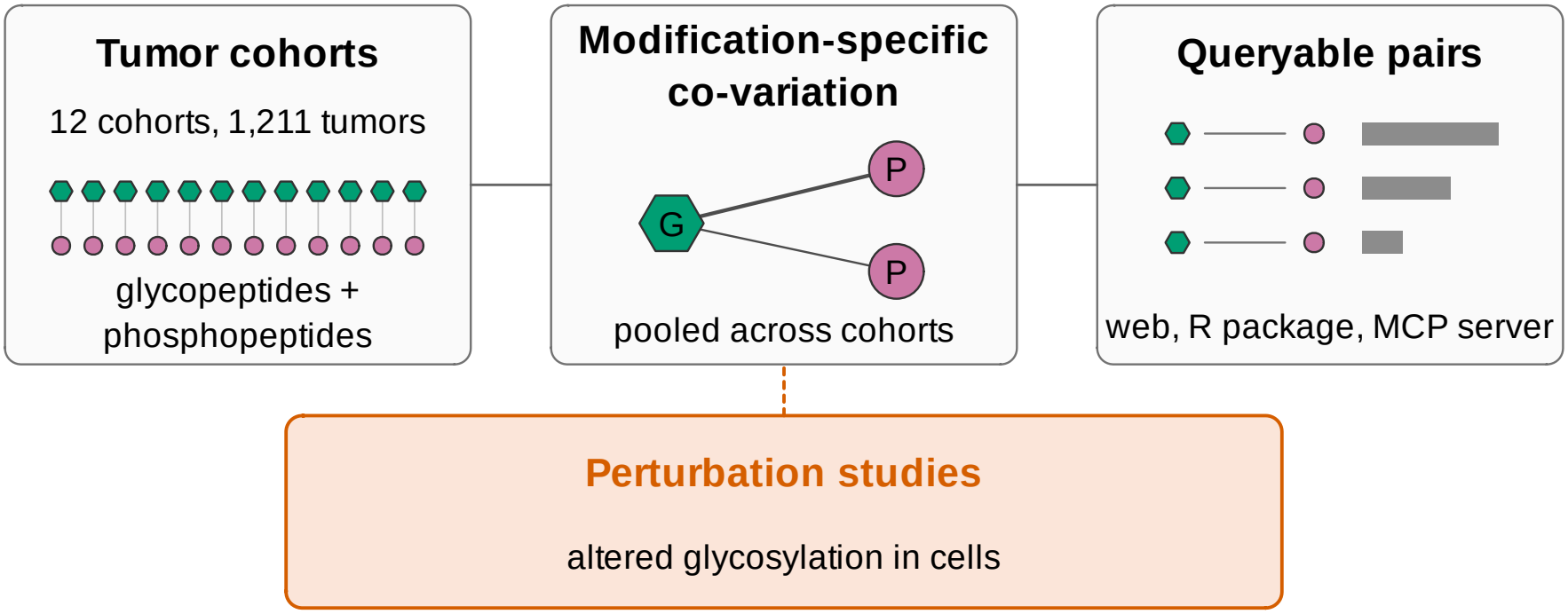

## Notes

https://ptmlink.biomarkercenter.org

### Competing Interest Statement

The authors have declared no competing interest.

