## Supplementary Materials for "PTMlink: a database resource for co-variation of glycosylation and phosphorylation in human tumors"

**Supplementary Tables** **7**

**Supplementary Figures** **10**

**Supplementary References** **11**

#### Supplementary Methods

##### S1 Data sources and sample handling

For the freeze cohorts, phosphosite abundances are the harmonized Baylor College of Medicine processing ($\log_2$ abundance), glycoform abundances the Johns Hopkins University intact-glycopeptide quantification (median-normalized $\log_2$ ratios), and protein abundances, used only in the annotations, the Baylor College of Medicine gene-level proteome. For the gastric and rare kidney cohorts, the site, glycopeptide and protein tables of the source studies were used.

Reference, quality-control and pooled channels were removed. In the gastric cohort, only tumors whose sample label occurred once in each matrix were used. The rare kidney study also re-quantified the clear-cell renal tumors of the freeze; only its own tumors were read, so no patient enters twice.

Non-finite phosphosite values were treated as missing. In the freeze glycopeptide matrices, a value that occurs three or more times within one sample column records a fixed bound rather than a measurement and was also treated as missing; zeros and rows duplicated across all columns were kept.

Supplementary Table S4 lists each input file with its sample-selection rule and SHA-256 checksum; patient-level matrices remain with their source repositories.

#### S2 Site keys and identifiers

Each glycopeptide was mapped to the canonical UniProt sequence of its protein, numbered from the initiator methionine, and assigned to the first complete sequon it contains; glycosites from peptides with several sequons are marked, since composition data cannot show which sequon carried the glycan. Glycopeptides assigned to more than one gene, lacking a complete sequon or lacking a parsable glycan composition were removed.

Freeze phosphosites keep the position of the harmonized processing (GENCODE v34); the canonical UniProt position, found from the site's sequence window, is reported beside it and forms the stable site identifier, and disagreements are marked. A phosphosite of the cohorts published later takes the key of the freeze phosphosite with the same gene and an identical fifteen-residue window; sites without a unique match were not used.

Where several peptides report the same glycoform in a cohort, the one quantified with the phosphosite in the most tumors represents it in that pair; ties between glycoforms for a glycosite's representative are broken by composition. Glycosites are annotated with sequon type, membrane topology and UniProt glycosylation evidence.

#### S3 Glycan classes

A composition counts HexNAc, hexose, fucose, NeuAc and NeuGc. Two HexNAc with at most three hexoses and no sialic acid is paucimannose; two HexNAc with four to nine hexoses and no other residue, oligomannose; and two HexNAc with ten to twelve hexoses and no other residue, glucosylated oligomannose. Three HexNAc with at least five hexoses, or four or more HexNAc with more hexoses than HexNAc plus one, is hybrid, and the other compositions with three or more HexNAc are complex.

Compositions without a two-HexNAc core, two-HexNAc compositions with fucose or sialic acid on four or more hexoses or with more than twelve hexoses, larger compositions with more fucoses than one core fucose plus one per antenna or more sialic acids than antennae, and NeuGc, which human tissue does not synthesize, are flagged. Composition does not resolve linkage, branching or fucose position, so a class names what the counts are consistent with.

#### S4 Perturbation studies

*Dose response.* The study's Spearman correlation between each glycopeptide and each EGF-regulated, dose-responsive phosphopeptide across the tunicamycin series (eight doses and a vehicle control) was converted to an exact two-sided $P$ value and corrected by BH, and pairs at $q < 0.05$ were kept, so this study observes the pair itself.

*Gene editing.* Glycoforms that differed between a knock-in or knock-out line and wild-type HEK293 cells were paired with localized phosphosites whose response to serum stimulation was specific to one line.

*Sialidase.* The glycoforms and localized phosphosites that the study reported as differing in either comparison, sialidase against control with and without EGF, were paired.

#### S5 Composition scores and the program-adjusted correlation

The eleven scores are the immune–stromal contrast (the ESTIMATE immune minus stromal score, orthogonalized to the ESTIMATE score) and proteomic scores for plasma proteins, proliferation, myeloid, lymphoid, erythroid, fibroblast and endothelial cells, unfolded-protein-response chaperones, the glycosylation machinery and secretory capacity. Each proteomic score is the median of standardized abundances over a fixed marker panel, omitting the genes of the site or pair being scored. The median uses the remaining measured markers, with no additional minimum panel fraction, and is missing if none is available. Supplementary Table S6 lists the marker genes of each panel, and Supplementary Tables S5 and S7 give the coverage of each covariate and each panel by cohort.

The program-adjusted correlation is computed in the cohorts with immune and stromal estimates and pooled as the main correlation is. A pair requires at least 20 tumors with every adjustment covariate and a weight $N - 3 - k_c$ of at least 12. The rare kidney cohort lacks the immune–stromal contrast and has no program-adjusted estimate.

#### S6 Held-out cohorts and normal adjacent tissue

*Held-out cohorts.* Intervals come from 2,000 bootstrap resamples of glycoproteins. The slope is read like the calibration slope of a prediction model (1); above one, the held-out correlations are stronger than the pooled ones. The two largest slope estimates, 1.20 and 1.02 in GC and PDAC, were near or above one; these are also the two cohorts in which the most pairs stand out (Figure 3A of the article).

*Slope and cohort size.* The slope was compared with two other measures of reproduction: the Pearson correlation between the held-out and pooled correlations of the selected pairs, and the share of those pairs whose held-out correlation has the pooled sign and a two-sided $P < 0.05$ in the cohort's own test. The slope depended less on cohort size: its Spearman correlation with the cohort's number of tumors was 0.28, against 0.51 for the Pearson correlation and 0.51 for the share of significant pairs.

The reason is that sampling noise in the held-out correlations, which grows as a cohort gets smaller, widens the interval of the least-squares slope without changing its expectation, given the selected pooled correlations. The Pearson correlation, whose denominator includes that noise, and the share of significant pairs both fall as a cohort gets smaller. The pooled correlations still carry their own uncertainty and selection, and cohorts can differ in true effects, eligible pairs and measurement precision, so the slopes are not shown to be independent of sample size.

*Normal adjacent tissue (NAT).* Each NAT cohort was first checked by permuting its phosphosite sample labels: across twenty permutations of 50,000 pairs, the share at $P < 0.05$ had to lie between 0.03 and 0.07. Agreement with the tumors was accepted when the lower 95 % bounds exceeded one half for the sign agreement and zero for the Spearman correlation; intervals come from 1,000 bootstrap resamples of glycosites. Tumors and NAT share patients and isotope-labeling sets, which may contribute, with shared biology, to the above-chance agreement of the non-significant pairs. The comparison therefore measures concordance and is not an independent replication.

#### S7 Known interactions and pathways

A STRING edge is one scoring at least 400 in the experimental or curated-database channel. Each gene pair was placed in the evidence bin of its strongest site pair. The strata were quintiles of the absolute parent-protein correlation, tertiles of each gene's STRING degree in the experimental and curated-database channels, and halves of its text-mining degree. The reference set is the gene pairs of a random 3 % sample of all scored site pairs. Stratum counts were combined as a Mantel–Haenszel risk ratio with Greenland–Robins 95 % intervals. The edges of the evidence bins were fixed before this release. Supplementary Figure S2B–C repeat the comparison with the gene-pair ranking in place of the evidence bins and a STRING edge defined by a combined score of at least 700.

#### S8 Literature relations

*Curation.* Relations were collected in rounds: a seed set of papers read in full; searches of PubMed titles and abstracts and of Europe PMC full texts, organized by protein family, for manipulations of *N*-glycosylation (sequon mutation, glycosylation enzymes, tunicamycin, NGI-1, sialidase, PNGase F) together with phosphorylation readouts; and a later full-text search in which each paper was read twice independently and disagreements were adjudicated. A paper was included when it was primary research, altered the *N*-linked glycosylation of a named glycoprotein, reported a phosphorylation readout of a named protein and stated a direction, recorded with a verbatim quotation. At merge, every PubMed identifier was checked and both genes were mapped to human UniProt accessions. Papers from the authors' laboratory, including the three perturbation studies, and papers whose data come from CPTAC tumor cohorts were excluded. Searches and extraction after the seed set were carried out by language-model agents (Claude, Anthropic) working from written briefs, with every query, decision and rejected paper recorded.

*Relations and matching.* A relation counts four kinds of manipulation of *N*-glycosylation: loss of a sequon (by mutation or deletion), enzymatic glycan removal, gain or loss of a glycosylation enzyme, and inhibitors. Any scored glycosite–phosphosite pair of the two proteins may support a relation. A relation $G$–$P$ is compared with gene pairs $G'$–$P$ that share its phosphoprotein, are not curated relations and have a site pair measured in at least two cohorts. $G'$ must match $G$ exactly in which of the perturbation studies that assayed $P$ also assayed its glycosites and in the binned number of glycosites they assayed, and lie within 0.25 of $G$ in $\log_{10}(1 + \text{PubMed records})$ and within 0.5 in the $\log_{10}$ number of its site pairs with $P$ measured in at least two cohorts; the nearest ten are kept, the assay bins are coarsened when fewer than five qualify, and a relation with fewer than five matches is left out. The expected count is the sum, over relations, of the share of each relation's matched pairs with perturbation-only support. Resampling and matched-control uncertainty are conditional on the fixed literature list and this matching procedure. Supplementary Tables S10–S13 give the literature relations, their comparison with the matched pairs, the matched pairs and the settings.

*Random rounds.* In each of 500 rounds, every relation was replaced by one of its own matched pairs, drawn at random, and the ratio was recomputed with each replacement compared with the relation's other matched pairs. The plan required that the pass rule, a lower 95 % bound above one, be met in at most 5 % of these rounds before the result was interpreted; it was met in 1.2 % of them. The median ratio over the rounds was 1.00 and the largest 2.12; none reached the observed 2.18.

#### S9 Tumor specificity

For a pair measured in cohort $c$ and in at least two other cohorts, with the weights $w_j = N_j - 3 - k_j$ and Fisher scores $z_j$ of the pooled test,

$$\bar{z}_{-c} = \frac{\sum_{j \neq c} w_j z_j}{W_{-c}}, \qquad W_{-c} = \sum_{j \neq c} w_j, \qquad z_{\Delta,c} = \frac{z_c - \bar{z}_{-c}}{\sqrt{1/w_c + 1/W_{-c}}}, \tag{1}$$

and BH correction of the two-sided $P$ values of $z_{\Delta,c}$ over all such pairs of cohort $c$ gives $q_\Delta$. The cohort's correlation counts as stronger than the background when $|\rho_c|$ exceeds $|\tanh \bar{z}_{-c}|$.

To check that $z_\Delta$ follows its null distribution, each cohort's tumors were split twenty times into random halves by patient and $z_\Delta$ was computed between the halves over the pairs, among 50,000 drawn, measured in at least 20 tumors of both halves; no pair was reported unless, in every cohort, the standard deviation of $z_\Delta$ over the splits lay between 0.9 and 1.1 and the share at $P < 0.05$ between 0.03 and 0.07. The standard deviation was 1.00 to 1.02 in every cohort, as expected when there is no difference. The difference that a change of phosphoproteomic pipeline alone produces was measured as $z_\Delta$ between two quantifications of the same 103 clear-cell renal tumors; its standard deviation was only 0.24.

The comparison at equal size was specified before it was run. A random 200,000 pairs (of a random set of genes) eligible in each cohort and measured there in at least 30 tumors were drawn. Each cohort's correlation was recomputed on 30 tumors drawn at random from the pair's own tumors, ten times, and compared with the unchanged background of the other cohorts, with BH within the cohort. The pairs of one draw share, as far as possible, the same randomly drawn tumors, which is why a cohort's draws differ. The criterion for a difference beyond size was that the cohort with the largest mean share had at least three times the median cohort's share and that its lowest draw lay above the highest draw of the cohort ranked seventh. GC met both conditions (mean 0.71 % against a median of 0.023 %; 8.4 % when the same pairs were tested on all of their tumors). PDAC had the second-largest share, 0.51 % (23 times the median); the criterion covers only the cohort with the largest share. Besides tissue composition and tumor biology, a pair that stands out in one cohort can reflect the measurement depth or the quantification pipeline of its source study (Section S1).

#### S10 The pooled test and its calibration

The pooled test combines the cohorts' Student tests by a weighted Stouffer method. With the correlations $\rho_c$, sample sizes $N_c$, covariate counts $k_c$ and weights $w_c$ of the article,

$$t_c = \rho_c \sqrt{\frac{N_c - 2 - k_c}{1 - \rho_c^2}}, \qquad u_c = \operatorname{sign}(\rho_c)\, \Phi^{-1}\{1 - F_{N_c-2-k_c}(-|t_c|)\}, \tag{2}$$

$$U = \frac{\sum_c \sqrt{w_c}\, u_c}{\sqrt{\sum_c w_c}}, \qquad \text{evidence} = -\log_{10}\{2\,\Phi(-|U|)\}, \tag{3}$$

where $F_\nu$ is the Student distribution function with $\nu$ degrees of freedom and $\Phi$ the standard normal distribution function. Approximate 95 % intervals of the pooled correlation are ±1.96 standard errors, $(\sum_c w_c)^{-1/2}$, on the Fisher scale, transformed back; for rank correlations they are slightly narrow (2).

In simulation, one million pairs without co-variation were drawn at each number of cohorts from one to twelve, each cohort with its own covariates and a sample size between 20 and its number of tumors (Supplementary Figure S1A). On 100 measured site pairs, an equal number in each of five bands of cohort count, the ranked residuals were permuted within each cohort's own samples and covariates (5,000 draws per pair) and the pooled test of the article recomputed; it rejected at 0.0502 to 0.0506 at a nominal 0.05 and at no more than 0.0011 at 0.001. These checks assess pointwise calibration in the settings examined, not the realized false-discovery fraction across the dependent family of site-pair tests. The additional $q$ value displayed among perturbation-supported pairs is corrected separately within that subset and changes neither the primary $q$ value nor the ranking.

#### S11 Software and release

The build is written in R (version 4.5.0; data.table 1.17.2, readr 2.1.5, dplyr 1.1.4, matrixStats 1.5.0 and arrow 20.0.0) and shell; figures were drawn with ggplot2 3.5.2 and patchwork 1.3.0. Each release ships a version record of the cohort data sources, the reference resources with their versions, retrieval dates and checksums, the software versions and the checksum of every scored table. Every number in this document and in the article is generated by script from the analysis tables of the build behind the release. Supplementary Tables S4–S15 are the sheets of one Excel file. Its first sheet lists them and its last sheet defines every column:

- S4: every input file, with its sample-selection rule and checksum;
- S5–S7: the coverage of the covariates and composition scores, and the marker genes of each score;
- S8–S9: the sources and each cohort's estimate for the pairs of Supplementary Table S2;
- S10–S13: the literature comparison of Section S8;
- S14: the candidate pairs that carry perturbation support;
- S15: the version record of release v1.0.0.

The complete candidate screen and the R script that reproduces it from the full data release are deposited on Zenodo. The website uses AlphaFold models (3) and SNFG glycan symbols (4).

### Supplementary Tables

**Supplementary Table S1.** Resources relating protein modifications, as described in each resource's publication and documentation. Matched profiles, glycosylation and phosphorylation measured in the same samples. PTMlink (shaded) is, to our knowledge, the first to provide a queryable score for site-specific relations between glycosylation and phosphorylation across tumor cohorts. PTM, post-translational modification; FDR, false discovery rate; TCGA, The Cancer Genome Atlas; CPTAC, Clinical Proteomic Tumor Analysis Consortium.

| Resource | Relates | Matched profiles | Resolution | Material | Association statistics |
|---|---|---|---|---|---|
| PhosphoSitePlus (5) | sites and their evidence | no | site | curated literature, proteomic data sets | none |
| GlyGen (6) | glycosylation sites, glycans, glycoproteins | no | site, glycan | integrated databases | none |
| GlyConnect (7) | site-specific glycan structures | no | glycoform | curated glycoproteomic studies | none |
| iPTMnet (8) | sites, enzyme–substrate relations, PTM-dependent interactions | no | site | text mining, curated databases | evidence-confidence score per record; no *P* |
| dbPTM (9) | sites, enzyme–substrate relations, tumor–normal phosphosite changes | no | site | curated literature, proteomic data sets | *P* for tumor–normal changes; no glycosite–phosphosite association *P* |
| PTMcode v2 (10) | site pairs within and between proteins | no | site pair | co-evolution, structural proximity, curation | co-evolution and conservation scores; no pooled tumor co-variation *P* |
| LinkedOmics (11) | feature–feature associations within a cohort; *P* values combined across cohorts | yes | gene, phosphosite; glycosite in ovarian cancer | tumor and normal tissue (TCGA, CPTAC) | *P* and FDR per association |
| Proteomic Data Commons (12) | data sets | yes | matrix | tumor and normal tissue (CPTAC and other programs) | none |
| Glycoprotein-Notebook (13) | glycoproteomic features across cancers; glycopeptide–phosphopeptide correlation in one cohort | yes | site, glycoform | tumor and normal adjacent tissue (CPTAC) | Spearman *P* per pair in its analysis code (one cohort); no combined score across cohorts |
| **PTMlink** | **glycosite × phosphosite pairs** | **yes** | **site, glycoform** | **tumor tissue (twelve cohorts) and normal adjacent tissue (six cohorts); perturbation evidence** | **queryable score per pair: pooled *P*, *q*, $I^2$ and intervals** |

**Supplementary Table S2.** Illustrative published protein relations and untested candidates: pooled estimates. $k$, cohorts; $N$, tumors; $\rho$, pooled partial Spearman correlation (95 % CI); Evid., $-\log_{10} P$; $q$, BH $q$ value; $I^2$, heterogeneity between cohorts; $\rho_{\text{prot}}$, parent-protein correlation; Adjusted CI, program-adjusted 95 % interval. The rows are illustrations, not a ranked list.

| Pair | $k$ | $N$ | $\rho$ (95 % CI) | Evid. | $q$ | $I^2$ | $\rho_{\text{prot}}$ | Adjusted CI |
|---|---|---|---|---|---|---|---|---|
| *Published protein relations* | | | | | | | | |
| ITGB4 N327 × AKT1 S124 | 4 | 247 | 0.38 (0.27, 0.49) | 8.87 | $3.4 \times 10^{-7}$ | 0.43 | 0.03 | (0.29, 0.51) |
| EGFR N603 × ALG3 S13 | 5 | 437 | 0.22 (0.13, 0.31) | 5.18 | $3.8 \times 10^{-4}$ | 0.60 | 0.03 | (0.11, 0.29) |
| ITGAV N615 × AKT1 S124 | 11 | 768 | 0.29 (0.22, 0.35) | 13.95 | $1.3 \times 10^{-11}$ | 0.81 | 0.19 | (0.19, 0.33) |
| ITGB1 N212 × PXN S533 | 9 | 771 | 0.26 (0.19, 0.32) | 11.73 | $1.2 \times 10^{-9}$ | 0.67 | 0.27 | (0.13, 0.28) |
| CD274 N192 × STAT1 S532 | 2 | 128 | 0.57 (0.43, 0.68) | 11.09 | $4.2 \times 10^{-9}$ | 0.20 | 0.34 | (0.03, 0.38) |
| CD44 N57 × MAPK3 T202 | 6 | 463 | −0.27 (−0.35, −0.18) | 7.76 | $2.9 \times 10^{-6}$ | 0.67 | −0.14 | (−0.25, −0.07) |
| *Untested candidates* | | | | | | | | |
| ADGRE5 N453 × NFRKB S298 | 8 | 550 | 0.36 (0.29, 0.43) | 16.61 | $5.5 \times 10^{-14}$ | 0.69 | 0.00 | (0.21, 0.38) |
| LRP1 N729 × DIDO1 S1456 | 8 | 735 | 0.30 (0.23, 0.36) | 14.83 | $2.1 \times 10^{-12}$ | 0.71 | −0.03 | (0.15, 0.29) |
| CD109 N41 × ATF2 S112 | 7 | 591 | 0.30 (0.22, 0.37) | 12.01 | $6.7 \times 10^{-10}$ | 0.85 | −0.04 | (0.15, 0.32) |
| CLN5 N203 × AHNAK S4908 | 11 | 927 | −0.23 (−0.29, −0.16) | 11.03 | $4.8 \times 10^{-9}$ | 0.40 | −0.01 | (−0.22, −0.08) |
| HYOU1 N596 × LARP7 S298 | 12 | 942 | 0.22 (0.15, 0.28) | 10.06 | $3.3 \times 10^{-8}$ | 0.62 | 0.10 | (0.04, 0.17) |
| IL6ST N131 × TMX1 S270 | 7 | 469 | 0.23 (0.14, 0.31) | 5.90 | $1.0 \times 10^{-4}$ | 0.15 | −0.09 | (0.09, 0.28) |

*Notes.* Published status concerns the protein relation, not experimental validation of the listed residue pair or the direction of its tumor co-variation. Sources: ITGB4–AKT1 (14, 15); EGFR–ALG3 (16); ITGAV–AKT1 (17); ITGB1–PXN (18); CD274–STAT1 (19); CD44–MAPK3 (20). ITGB4–AKT1 and EGFR–ALG3 are the worked examples of the article; the other four published relations were chosen after inspecting the release scores, each as the highest-evidence reported pair of its protein relation measured in at least two cohorts; the candidates are pairs from a curated list that pass the complete screen. Background on the candidate proteins: ADGRE5 (CD97) (21); NFRKB, a subunit of the INO80 complex (22); LRP1 (23); DIDO1 (24); CD109 (25, 26); ATF2 (27); CLN5 (28–30); AHNAK (31, 32); HYOU1 (ORP150) (33); LARP7 (34); gp130 (IL6ST), whose glycosylated extracellular domains are disulfide-bonded and whose N131 is N109 of the mature protein (35), and TMX1, which works with the lectin chaperone calnexin (36). Earlier studies identified the glycosites of EGFR (37) and, by glycoproteomics, ADGRE5 N453 (38), LRP1 N729 (38–40) and HYOU1 N596 (38, 41).

**Supplementary Table S3.** Literature relations with perturbation support and no significant tumor co-variation. Manipulation, how the study altered glycosylation (an enzyme change may be shown for the pathway rather than for the named glycoprotein; a sialidase treats whole cells, and the glycoprotein named is the one assayed); System, where phosphorylation was measured. An ERK1/2 immunoblot is recorded as MAPK3. ISC, intestinal stem cell; Siglec-7+, expressing sialic-acid-binding immunoglobulin-like lectin 7; NRP1-DeltaE4 and -DeltaE5, splice variants lacking exon 4 or 5.

| Glycoprotein | Phosphoprotein | Study | Manipulation | System |
|---|---|---|---|---|
| CD151 | STAT3 | Zhang 2026 (42) | enzymatic removal | Human HTR-8/SVneo extravillous trophoblast cells cocultured with Siglec-7+ NK-92MI cells |
| CD276 | MTOR | Wang 2023 (43) | FUT8 inhibitor FDW028 | Human colorectal cancer cell lines SW480 and HCT-8 |
| CD276 | STAT3 | Zhang 2026 (42) | enzymatic removal | Human HTR-8/SVneo extravillous trophoblast cells cocultured with Siglec-7+ NK-92MI cells |
| CDH2 | MAPK3 | Noor 2021 (44) | enzyme gain or loss | Human HEK293T cells, POMGNT1 knockout versus wild type |
| FGFR1 | MAPK3 | Cazet 2014 (45) | enzyme gain or loss | Human glioma SKMG-3 cells |
| ITGA5 | EGFR | Hang 2015 (46); Hang 2016 (47) | sequon mutant | Human HeLa (integrin $\alpha$5 knockout, reconstituted with $\alpha$5 mutants), U-251MG and MDA-MB-231 cells |
| ITGA5 | MAPK3 | Hang 2016 (47) | sequon mutant | Hamster CHO-B2/EGFR cells reconstituted with integrin $\alpha$5 variants |
| ITGAV | MAPK3 | Chiodelli 2012 (48) | enzymatic removal | Transformed fetal bovine aortic endothelial GM7373 cells |
| ITGB3 | MAPK3 | Chiodelli 2012 (48) | enzymatic removal | Transformed fetal bovine aortic endothelial GM7373 cells |
| MUC16 | MAPK3 | Rao 2015 (49) | sequon mutant | Mouse NIH-3T3 fibroblasts stably transfected with human MUC16 constructs |
| NCAM1 | MAPK3 | Gong 2017 (50) | enzyme gain or loss | Human small-cell lung cancer H446 cells |
| NPC1 | MTOR | Duan 2024 (51) | enzyme gain or loss, sequon mutant | Mouse ileal crypts, ISC-specific Fut2 knockout versus wild type; mouse ileal organoids, Npc1 N961Q |
| NRP1 | PTK2 | Huang 2019 (52) | sequon loss (splice variants) | Human colorectal cancer HCT116 cells stably expressing NRP1-WT, NRP1-DeltaE4 or NRP1-DeltaE5 |

### Supplementary Figures

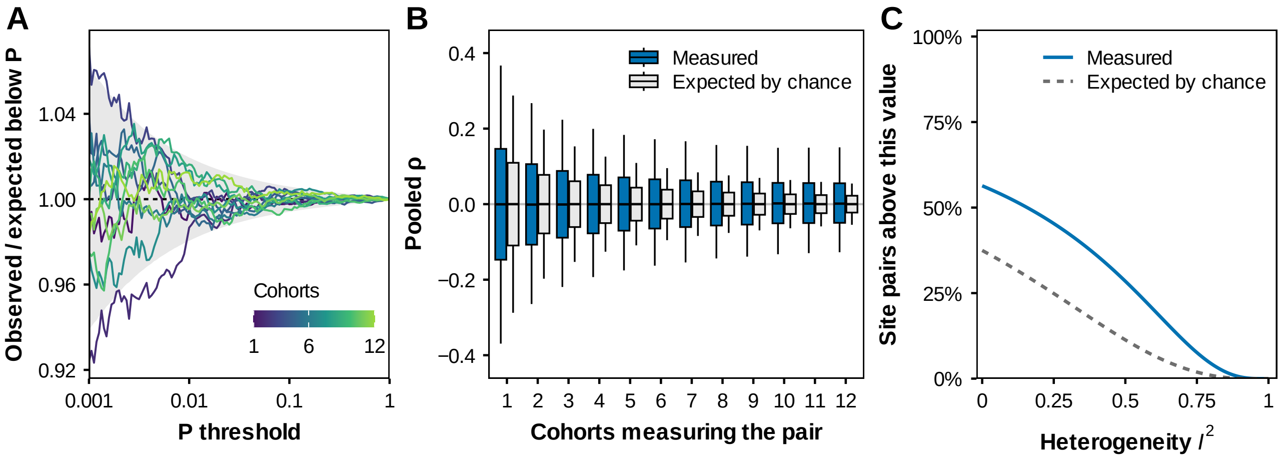

**Supplementary Figure S1.** Quality control of the pooled statistic. (**A**) Observed over expected number of simulated pairs without co-variation below each $P$ threshold, by number of cohorts; shading, 95 % range from counting noise. (**B**) Pooled correlations of all scored site pairs by number of cohorts, measured and expected without co-variation; boxes, quartiles; whiskers, 5th and 95th percentiles. (**C**) Share of site pairs measured in at least two cohorts whose $I^2$ exceeds each value, measured and expected when the cohorts share one correlation.

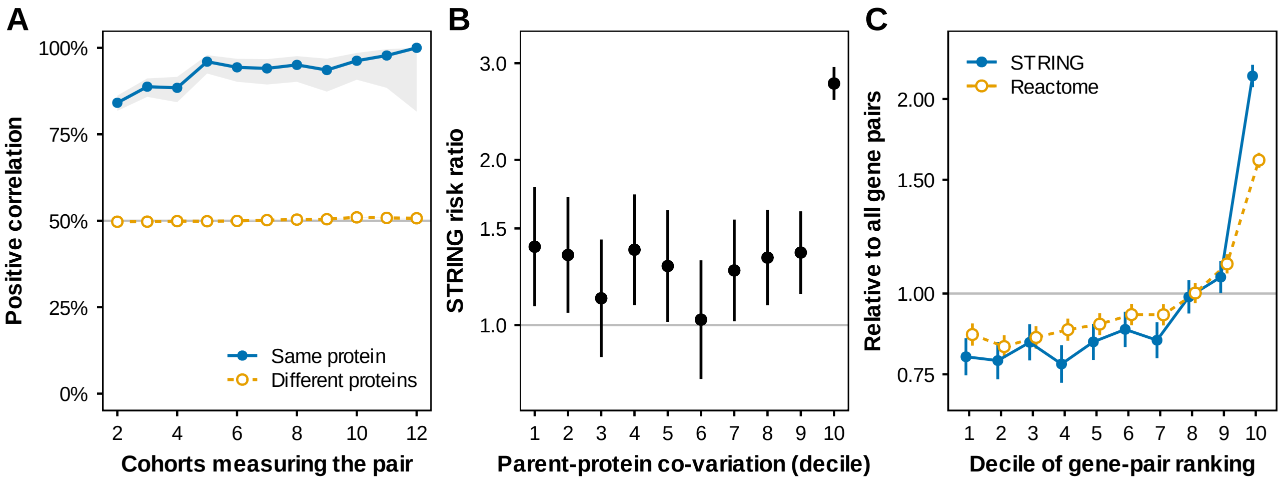

**Supplementary Figure S2.** Shared protein abundance and known interactions. (**A**) Share of pairs with a positive correlation by number of cohorts, for two sites on the same protein (filled, solid line) or on different proteins (open, dashed line); shading, Wilson 95 % intervals. (**B**) Within each decile of parent-protein co-variation evidence, the share of gene pairs with a STRING edge in the top tenth of the gene-pair ranking over that share in the rest; bars, 95 % intervals. (**C**) Gene pairs with a STRING edge or Reactome co-membership in each decile of the gene-pair ranking, relative to all gene pairs; bars, 95 % intervals. The tenth decile is the strongest (Section S7).
